# From scales to armour: scale losses and trunk bony plate gains in ray-finned fishes

**DOI:** 10.1101/2020.09.09.288886

**Authors:** Alexandre Lemopoulos, Juan I. Montoya-Burgos

## Abstract

Actinopterygians (ray-finned fishes) are the most diversified group of vertebrates and are characterized by a variety of protective structures covering their tegument, the evolution of which has intrigued biologists for decades. Paleontological records showed that the first mineralized vertebrate skeleton was composed of dermal bony plates covering the body, including odontogenic and skeletogenic components. Later in evolution, the exoskeleton of actinopterygian’s trunk was composed of scale structures. Although scales are nowadays a widespread tegument cover, some contemporary lineages do not have scales but bony plates covering their trunk, whereas other lineages are devoid of any such structures. To understand the evolution of the tegument coverage and particularly the transition between different structures, we investigated the pattern of scale loss events along actinopterygian evolution and addressed the functional relationship between the scaleless phenotype and the ecology of fishes. Furthermore, we examined whether the emergence of trunk bony plates was dependent over the presence or absence of scales. To this aim, we used two recently published actinopterygian phylogenies, one including > 11,000 species, and by using stochastic mapping and Bayesian methods, we inferred scale loss events and trunk bony plate acquisitions. Our results reveal that a scaled tegument is the most frequent state in actinopterygians, but multiple independent scale loss events occurred along their phylogeny with essentially no scale re-acquisition. Based on linear mixed models, we found evidence supporting that after a scale loss event, fishes tend to change their ecology and adopt a benthic lifestyle. Furthermore, we show that trunk bony plates appeared independently multiple times along the phylogeny. By using fitted likelihood models for character evolution, we show that trunk bony plate acquisitions were dependent over a previous scale loss event. Overall, our findings support the hypothesis that tegument cover is a key evolutionary trait underlying actinopterygian radiation.

**Impact Summary:** Ray-finned fishes (actinopterygians) are the most diverse vertebrate group in the world. The majority of these fishes possess scales as a protective shield covering their trunk. However, several lineages display a body armour composed of trunk bony plates or are devoid of any protective structures. The diversity and the transitions between different tegument coverage types have not been previously studied in an evolutionary framework. Here, we investigate which structure was present at the origin of ray-finned fishes and how the different phenotypes emerged through time.

We show that a scaled tegument was the most widespread sate along ray-finned fish evolution, yet scale losses occurred multiple independent times, while acquiring scales again almost never happened. Moreover, we reveal that scaleless teguments most probably led species to change their ecology and colonise the floors of oceans and water bodies. The functional advantages of a scaleless tegument in a benthic environment are yet to be demonstrated, but the increased cutaneous respiration could be an explanation. We show that trunk bony plates also emerged independently multiple times along the evolution of ray-finned fishes but these armours protecting the trunk can only appear after a scale loss event. Therefore, while the acquisitions of trunk bony plates are phylogenetically independent, they need a “common ground” to emerge. All together, our findings provide evidence that the various tegument covers have contributed to the outstanding diversification of ray-finned fishes.

## Introduction

Ray-finned fishes (Actinopterygii) represent the most diversified vertebrate lineage in the world, with more than 33 thousand described species (Alfaro 2018). One of the most prominent features among Actinopterygii representatives is the presence of scales in their trunk tegument forming a protective layer. Scales can display various shapes and structures, as they can contain different compounds and differ in histological characteristics (Moyle and Cech 2004). The diversity of scales has created some confusion in the scientific community, because different skeletal elements have been referred as scales despite being of different origin (Schultze 2018). Yet, given the great diversity and the complexity of these structures, a consensus over their nomenclature and classification still needs to be established based on a comprehensive understanding of their evolutionary origin (Sire et al. 2009; Vickaryous and Sire 2009). In this study, we primarily focus on two categories of mineralized structures developing within the tegument of actinopterygians, micromeric scales and macromeric trunk bony plates.

Scales, as differentiated micromeric dermal skeletal elements (*sensu* Sire 2003 and Sire et al. 2009) were present in the ancestral lineage that gave rise to Actinopterygii and Sarcopterygii (Sire et al. 2009). Therefore, scales are considered a plesiomorphic trait for ray-finned fishes and today the majority of them possess some type of scales (Gemballa and Bartsch 2002; Sire et al. 2009). Based on different histological and morphological properties, scales have been classified in two main groups: ganoid scales (in Protopteridae (bichirs) and Lepisosteiformes (gars) [Meunier and Brito 2004; Ichiro et al. 2013]) and elasmoid scales (in the majority of actinopterygian lineages; e.g. Sire et al., 1997; Mongera and Nüsslein-Volhard, 2013). All scales possess a bony layer (e.g. bony-ridge, lammellar bone) in their structure (Benthon 2004; Moyle and Cech 2004; Zhu et al. 2012). Thus, scales are a bony structure covered with a scale-specific odontogenic-like tissue, in general. The nature of the odontogenic-like cover and the scale organization then define the type of scale (e.g. ganoin in ganoid scales; Ichiro et al. 2013). Therefore, two components are in general necessary for the formation of a scale: a) a bone micromeric structure; and b) an odontogenic-like cover tissue that is scale-specific (but this tissue is sometimes reduced or even absent).

Trunk bony plates (TBP) represent another type of tegument protection, which is present in some extant actinopterygians. The origin of TBP can be traced back to the first vertebrate mineralized skeleton, which was composed of TBP covered with an odontogenic tissue (Keating and Donoghue 2016). Independently of their evolutionary history, TBP *sensu lato*, can be differentiated from scales as macromeric tegumental elements composed of bone only (i.e. lacking the odontogenic-like cover). TBP, as macromeric tegument structures, reappeared in specific actinoptetygian lineages. For instance, the iconic seahorse (Syngnathidae) exoskeleton is made of dermal bony plates covering the entire body (Lees et al., 2012; Porter et al., 2013). Other examples are the Callichthyidae and the Loricariidae, two species-rich families of Neotropical catfishes, that have their trunks covered with TBP (Sire 1993; Covain et al. 2016; Rivera-Rivera and Montoya-Burgos 2017). Interestingly, micromeric scales and macromeric TBP seem to be mutually exclusive as no extant fish displays both exoskeletal structures in the trunk.

Despite the widespread occurrence of protective elements in the tegument of fishes, several lineages within actinopterygians display a naked skin, i.e. devoid of any scales or any other protective structures. Whether the lack of scales in several ray-finned fishes is a result of independent scale loss events rather than multiple independent appearances of scales has not been formally assessed. Nevertheless, the putative selective advantage of a scaleless skin is compelling. Some functional advantages have been suggested, such as an increased cutaneous respiration (Park and Kom 1999; Park 2002), or a relatively higher expression of immune genes after a parasitic infection as measured in scaled versus scaleless skin regions of salmons (Holm et al. 2017). Yet, the extent of the advantages and disadvantages of having a scaleless tegument is unclear. Nevertheless, we observed that scaleless fishes belonging to different lineages tend to have a benthic habitat preference. In addition, they present a similar overall morphology corresponding to the one typically found in bottom-dwelling species (e.g. inferior mouth, flattened abdomen or body) according to the classification of Moyle and Cech (2004). Whether a relationship between a scaleless tegument and habitat preference exists in ray-finned fishes needs to be examined further.

In this study, we investigated the drivers of trunk tegument evolution in actinopterygians. We first hypothesized that the loss of scales may be related to a bottom-dwelling lifestyle, as this state could result in functional advantages in a benthic environment. Second, as apparently scales on the skin cannot co-occur with TBP in a same fish species, we tested the hypothesis that the loss a scales is an evolutionary prerequisite for the re-emergence of TBP. To test these hypotheses, we inferred the evolutionary history of the emergence and disappearance of scales and TBP along the evolution of actinopterygians. To this aim, we used two recently published ray-finned fishes phylogeny, one containing 304 species (Hughes et al. 2018) and the other 11,638 species (Rabosky et al. 2018). The magnitude of this dataset allowed us to have a precise view on actinopterygian evolution. We collected data regarding habitat preference and trunk tegument characteristics for each species of the phylogenies. We then performed ancestral state reconstructions and we investigated the associations between traits using methods of linear regression for binary data and likelihood model fitting for character evolution.

## Material and Methods

### Phylogeny

To perform the ancestral state reconstruction analyses and to account for the possible effect of (i) variation in the phylogenetic inferences (phylogenetic uncertainty) and (ii) phylogenetic relatedness of the traits in the correlation analyses, we used two recently published ray-finned fishes phylogenies (Hughes et al. 2018; Rabosky et al. 2018).

Hughes et al. (2018) published a robust and well-resolved phylogeny obtained by using 1105 orthologous exons of 305 species representing all actinopterygian lineages, including most of the lineages displaying the traits examined in this study. One species, *Xenopus tropicalis*, used as an outgroup in Hughes et al. (2018) phylogeny was excluded from our analysis as it was irrelevant in the context of our study. Rabosky et al. (2018) phylogeny was reconstructed based on a 27 genes alignment for 11,638 species (with a substantial amount of missing data, see Rabosky et al. 2018). It is currently the most complete phylogeny as it contains almost all actinopterygian species.

### Tegument characteristics, morphology and habitat preference

Information about the traits displayed by fish species were collected in two books (Moyle and Cech 2004; Nelson et al. 2016) and in Fishbase (Froese and Pauly 2011). When information was lacking or unclear in these three main sources, species characteristics were extracted from the specialized literature. As the presence of scales is the ancestral trait of actinopterygians (Friedman and Brazeau 2010; Qu et al. 2013) we reported evidence for changes of traits, such as the absence of scales in the species or the presence of trunk bony plates (Table S1 for the 304 species dataset and Table S2 for the 11,638 species dataset).

To assess the link between absence of scales and habitat preference, we used as a baseline the classification of Moyle and Cech (2004) that links morphology to habitat preference. They described 10 different types of morphology-habitat associations, classified into five main categories. Out of these categories, four include fishes with middle or surface water habitat preference, while one category consists of fishes with bottom habitat preference (bottom-dwellers). According to Moyle and Cech (2004), this bottom-dwelling category contains five types of morphologies: 1) bottom-rovers (e.g. Siluriformes), 2) bottom-clingers (e.g. Cottidae), 3) bottom-hiders (e.g. some Percidae), 4) flatfish (e.g. Pleuronectiformes) and 5) rattail (e.g. Macrouridae). Here, we individually assessed and assigned each species present in the phylogenies to either the bottom-dwelling category or to the non-bottom-dwelling super-category. In addition, we used available literature for refining the species habitat preference in ambiguous cases. For instance, even though Moyle and Cech (2014) do not consider eel-like fish as bottom-dwellers, some eel-like species are bottom-associated such as swamp eels (*Synbranchus marmoratus*). The final corrected species allocation to the bottom-dwelling category and the non-bottom-dwelling super-category are presented in Table S1 for the 304 species dataset (Hughes et al. 2018) and in Table S2 for the 11,638 species dataset (Rabosky et al. 2018).

To differentiate TBP from scales we used the description by Sire and Huysseune (2003). Based on different phylogenetic, developmental and histological characters, they described 10 different dermal skeletal elements in fish trunks, which can be subdivided into (i) large macromeric bony plates and (ii) small micromeric scale-like elements. Trunk macromeric bony plates include cranial and postcranial dermal bones, and scutes (trunk bony plates specific to some Neotropical catfish), which we refer to as *TBP*. Trunk micromeric scale-like structures include odontodes (superficial structure with dental tissues), ganoid scales (of polypterids and lepisosteids) and elasmoid scales and they were here referred to as *scales*. In our study, we did not consider oral and extra-oral teeth or denticles as trunk scale-like elements as they represent more complex structures including dentine, enamel-like covers, a pulp cavity, a particular attachment to the underlying bone and an innervation in most cases. For each species, the tegument characteristics are presented in Table S1.

### Ancestral state reconstruction

We performed two different ancestral state reconstructions for the presence / absence of scales and for the presence / absence of TBP. We first used a stochastic mapping approach for morphological characters (Huelsenbeck et al., 2003). We used the make.simmap function in the phytools package v.06.99 (Revell 2012) in R environment v. 3.6.1. To infer for best model of transition rate, we compared AIC scores between the equal rate (ER) and all rates different (ARD) models (Table S3). The Q matrix for transition rates was sampled based on posterior probabilities after 250’000 generations (Q = “mcmc”) with a burnin phase of 10,000 generations. Prior probability distributions were set empirically with the option prior = use.empirical = true.

Second, we performed ancestral reconstructions using Bayestrait 2.0 (Pagel et al. 2004). For these reconstructions, we compared uniform and exponential reverse-jumping hyperprior (Pagel 2004). By comparing likelihood scores obtained by stepping stones (100 to 1000), we used the logBF factor to identify the best model for each scenario (Pagel et al. 2004; Table S3). To constrain jump acceptance rates for each model between 0.2 and 0.4, we used hyperpriors ranging from 0 to 30 as recommended by the software manual. We performed respectively 50,000,000 MCMC iterations for the 304 species phylogeny and 10,000,0000 iterations for the 11,638 species phylogeny. Trees and node probabilities were visualized using Treegraph 2 (Stöver and Müller 2010).

Finally, to evaluate how phylogenetic uncertainty could influence the ancestral reconstruction, in addition to working with two different datasets (Hughes et al. 2018; and Rabosky et al. 2018), we performed a multi-tree ancestral state reconstruction using the 304 species dataset of Hughes et al. (2018). We performed a phylogenetic inference with Exabayes (Aberer et al. 2014) based on the protein super-alignment provided by Hughes et al. (2018). We used the Hughes et al. (2018) best phylogeny as a starting tree in Exabayes and the other parameters were set to default. Iterations were executed until convergence was reached and visualised through Tracer v.1.7 (Rambaut et al. 2018). From the output of this analysis, we used a subset of 1000 trees to perform a multi-tree ancestral state reconstruction in Bayestrait 2.0. The parameters used were the same as the ones used on the single best tree analysis (Table S3).

### Association and dependency analyses

For the scaled/scaleless fish dataset and the presence/absence of TBP dataset, we calculated the *D* value, which is an index that indicates whether binary traits evolve independently or evolve according to the phylogeny under a Brownian motion model (Fritz and Purvis 2010). Thus, this value indicates to what extent the evolution of the traits is linked to the phylogeny (0 = no relationship; 1 = full dependency). We calculated this value using the phylo.D function in caper library v.1.0.1 (Orme et al. 2013).

To test the hypothesis that the scaleless phenotype is associated with a benthic habitat preference, we performed a linear regression for binary (discrete) data using the binaryPGLMM function in ape package v.5.3 (Paradis et al. 2004). Using this function, we tested whether the presence/absence of scales was explained by the habitat preference, and accounting for the phylogeny (Scale.State∼Ecology+[Phylogeny]). Parameters were set to default and convergence of the model was assessed using the build-in function.

To test the evolutionary relationship between scales and TBP, and more specifically whether the emergence of TBP was dependent over the absence of scales, we studied the association between presence/absence of TBP and presence/absence of scales using the fitPagel function in the phytools package (Revell 2012). This function is designed to analyse the coevolution of two traits and the way they are linked over the course of time by providing a phylogeny as an input to the method.

## Results

### Scale condition and habitat preference

Out of the 304 species considered in the phylogeny of Hughes et al. (2018), we identified 40 species as being scaleless (Table S1). 70 species were considered as bottom-dwellers, and 234 as non-bottom-dwellers. Ancestral state reconstructions were virtually the same with both stochastic mapping (Fig. 1 and Fig. S1) and Bayesian methods (Fig. S2 and Fig. S3). In both reconstructions, we identified 11 scale loss events The phylogenetic index *D* was not significant (Table S4) indicating that scale loss events bear no phylogenetic signal, that is, they occurred independently in different parts of the phylogeny. One event of scale re-acquisition following a loss was also inferred, namely in the *Anguilla* genus. However, this re-acquisition was observed only in the stochastic mapping reconstruction (Fig. 1 and Fig. S1), not in the Bayestrait reconstruction (Fig. S2 and Fig. S3). Results were similar when taking phylogenetic uncertainty into account by analysing a set of 1000 trees with Bayestrait (Fig. S3). However, an additional scale re-acquisition event was inferred with this reconstruction. This event occurred in the Opisthognathidae family (Fig. S3).

**Fig. 1.**
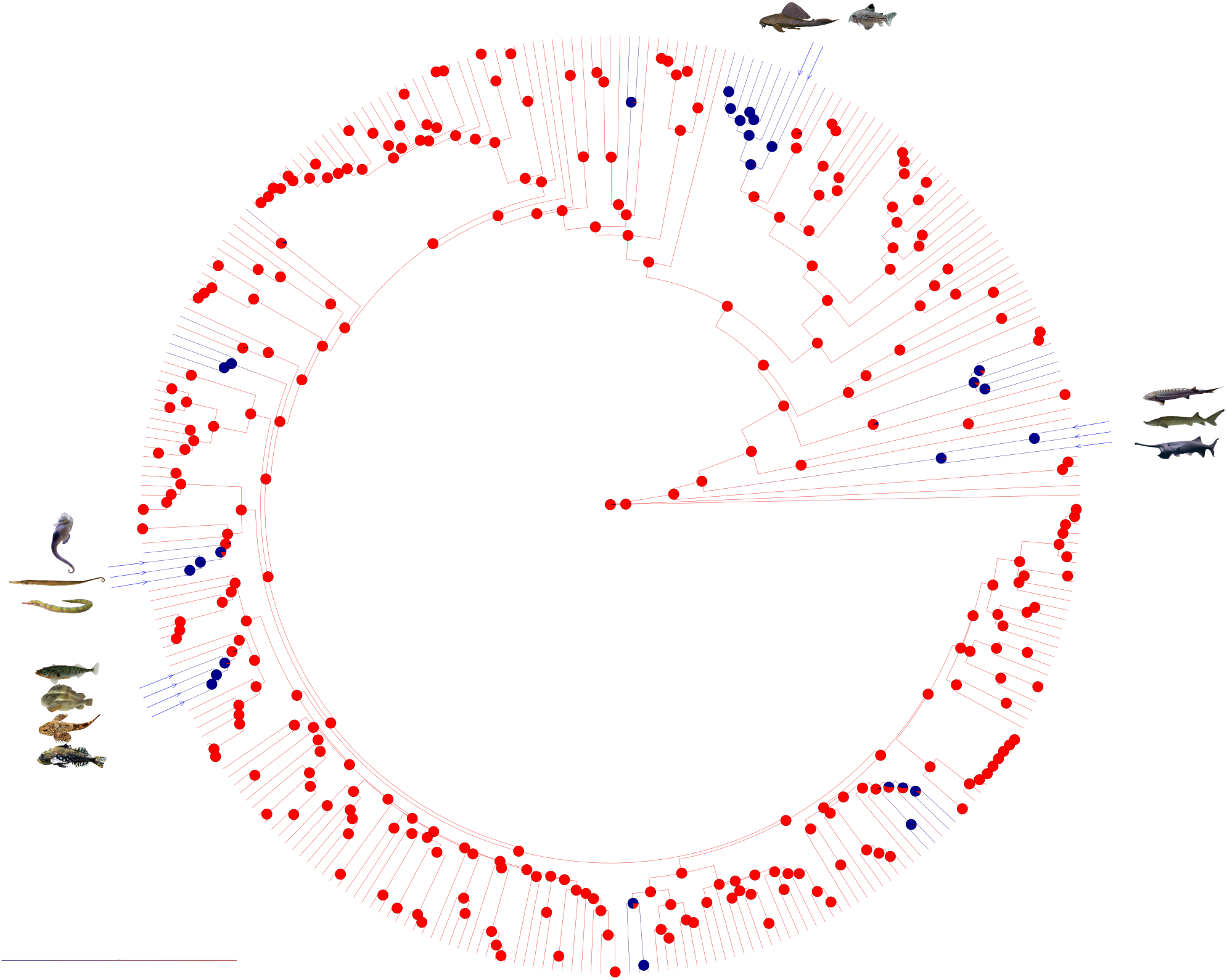
Reconstruction through stochastic mapping of the scale presence/absence on a phylogenetic tree of 304 Actinopterygii species (modified from Hughes et al., 2018, see Fig S1 for the detailed tree). Blue clades correspond to scaled taxa, while red color indicates lineages that underwent a scale loss event. Fish illustrations represent species displaying trunk dermal bony plates (TBP). The four distinct gains of TBP occured in distant lineages, yet always after a scale loss event. On the right, three species of Acipenseriformes: *Polyodon spathula, Acipenser sinensis, Acipenser naccarii;* on top, two species of Siluriformes: *Corydoras julii, Pterygoplichtys pardalis*; on the left three species of Syngnathiformes: *Syngnathoides biaculeatus, Syngnathus scovelli, Hippocampus erectus* and on the bottom left, four species of Gasterosteiformes: *Gasterosteus aculeatus, Cyclopterus lumpus, Cottus rhenanus, Myoxocephalus scorpius*.

In the 11,638 species dataset of the phylogeny by Rabosky et al. (2018), we identified 2,310 species as scaleless and 4,169 as bottom-dwelling. In the ancestral state reconstruction, 32 and 43 scale loss events were inferred with the stochastic mapping method (Fig. 2 and Fig. S4) and the Bayestrait method (Fig. S5). In contrast, 10 and 13 events of scale acquisition were inferred with the stochastic mapping and the Bayestrait methods, respectively. The phylogenetic index D was not significant, indicating that these trait changes are not phylogenetically linked (Table S4).

**Fig. 2.**
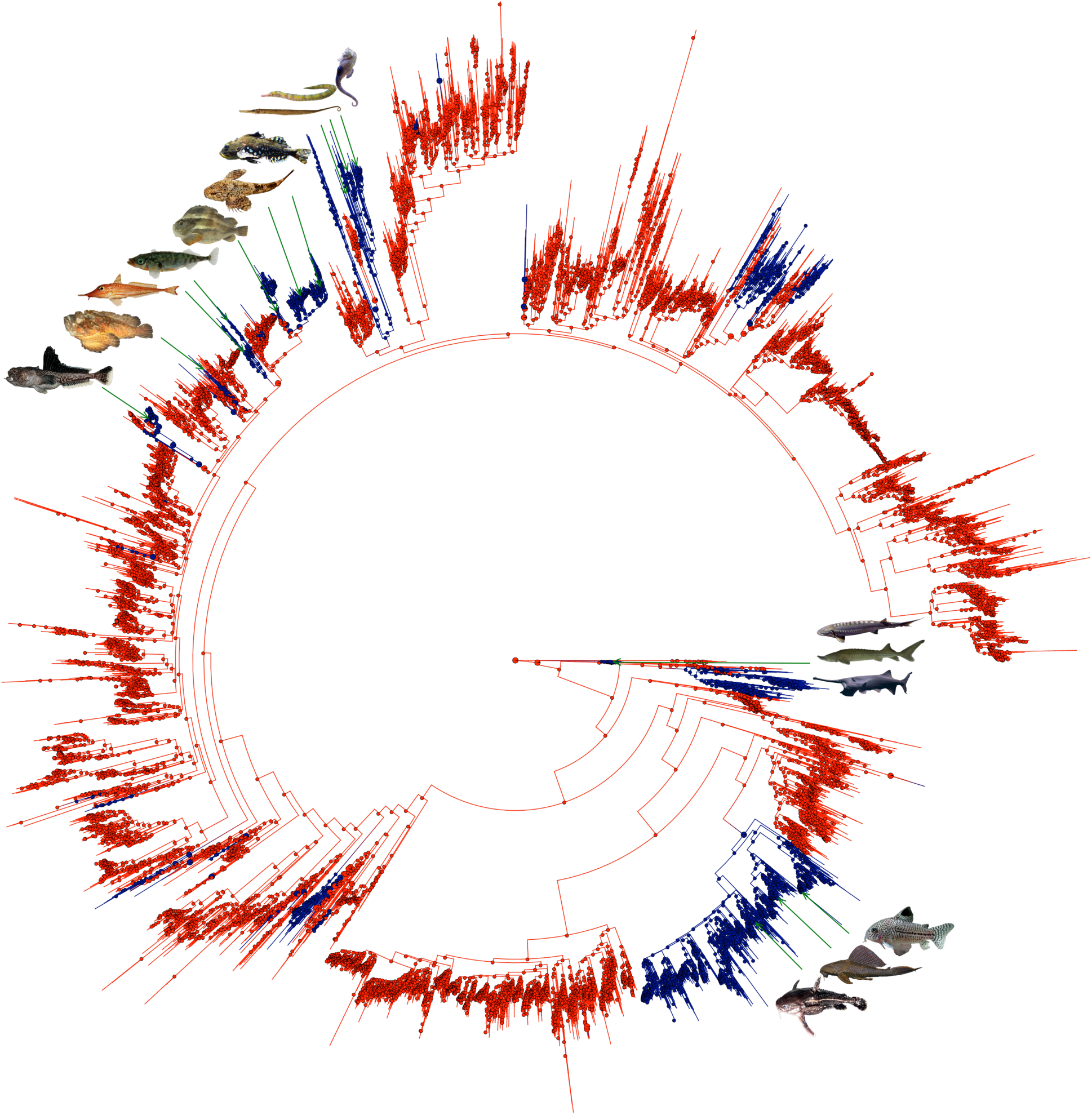
Ancestral trait reconstruction through stochastic mapping of the scale presence/absence on a phylogenetic tree of 11,638 Actinopterygii species (modified from Rabosky et al., 2018, see Fig S5 for the detailed tree). Blue clades indicate scaled taxa, while red color correspond to scaleless taxa. Pictures correspond to species representing lineages displaying TBP. These are found in several unrelated lineages, yet always after a scale loss event. Species illustrated in Fig. 1 are also represented here, in addition to other species not included in the dataset of Fig. 1. On the top left, ten fishes with TBP gains are illustrated. From left to right, one species of Perciformes: *Pogonophryne barsukovi* (1^st^ gain). Two species of Scorpaeniformes: *Synanceia verrucosa* and *Peristedion gracile* (2^nd^ and 3^rd^ gains). Within Gasterosteiformes, four species illustrate the fourth (*Gasterosteus aculeatus*) and the fifth TBP gain (*Cyclopterus lumpus, Cottus rhenanus, Myoxocephalus Scorpius*). Three species of Syngnathiformes: *Syngnathoides biaculeatus, Syngnathus scovelli, Hippocampus erectus* represent the seventh gain. On the center right, three species of Acipenseriformes: *Polyodon spathula, Acipenser sinensis, Acipenser naccarii* illustrate a distinct TBP gain event. Finally, on the bottom right, three species of Siluriformes: *Corydoras julii, Pterygoplichtys pardalis* and *Acanthodoras spinosissimus* are other examples of TBP gain.

The linear regression analyses to test for the association between the scaleless state and habitat preference showed with both datasets that the scaleless state and a benthic lifestyle are tightly linked (Table S5). Thus, fish presenting a scaleless tegument are potentially more likely to display a benthic habitat preference.

### Trunk bony plates emerge on a scaleless tegument

In the 304 species dataset, we identified 12 species displaying TBP, distributed over nine families. In both ancestral state reconstructions of presence/absence of TBP, stochastic mapping (Fig. S6) and Bayestrait (Fig. S7), 4 events of TBP acquisition were inferred. In contrast, no loss of TBP was inferred. The phylogenetic index *D* was not significant (Table S4), meaning that TBP appeared independently in different parts of the phylogeny. Results were almost identical when taking phylogenetic uncertainty into account by using a set of 1000 trees (Fig. S8). Only one additional TBP gain was identified in the Siluriformes order, but with a poor probability support.

In the 11,638 species dataset, 823 species displayed TBP (Table S2). The phylogenetic index *D* was also not significant (Table S4) for plate appearance. In the stochastic mapping reconstruction (Fig. S9), 16 plate gains and 6 plate losses were identified, while 21 plate gains and 11 plate losses were identified in the Bayestrait reconstruction (Fig. S10).

We assessed whether TBP appearance depends on a specific tegument condition using the fitPagel test. The results indicate that for both the 304 and 11’638 fish species datasets, the presence of TBP is dependent on a specific tegument scaling state (Table S5). More precisely, TBP have a significant and strong tendency to appear in a scaleless tegument.

## Discussion

Ray-finned fishes form the most species-rich group of extant vertebrates, and the reasons of their evolutionary radiation remains unclear. A number of functional innovations have been put forward to explain the wide radiation in Acanthomorpha, the main actinopterygian subgroup (Wainwright and Longo 2017), but the protection provided by their scaled exoskeleton is often forgotten. Yet, material engineers have demonstrated the mechanical and protective properties of fish scales (e.g. Zu et al. 2012), providing empirical evidence supporting that scales could be another functional innovation explaining the radiation of ray-finned fishes in the aquatic environment.

### Absence of scales is associated to habitat preference

Although a scaled tegument is one of the main characteristics of ray-finned fishes, the loss of scales occurred several times during the evolution of this group. Interestingly, we found that the presence/absence of scales bear virtually no phylogenetic signal indicating that a scaleless state arose independently in distant fish orders, as for instance in the Siluriformes or in the Acipenseriformes (Fig. 1 and Fig. 2 for the 304 and 11,638 species datasets, respectively).

Interestingly, our results based on both datasets revealed a tight association between the scaleless state and a benthic way of life. However, the strong correlation we revealed does not indicate whether the scaleless phenotype is a cause or a consequence of a benthic habitat preference. If scale loss were a consequence of a benthic ecology, then we would expect virtually no open water species displaying a scaleless tegument. To the contrary, if a benthic habitat preference were a consequence of scale loss in ancestors with open water habitat preference, then we would expect at least some scaleless taxa in open waters, as the loss of scales would initially occur there, before any possible translocation into the benthic habitat. After a careful examination of our table of the scaling status and the habitat preference, it appears that 19 scaleless fish families live in open waters (Table S6). We can mention, for instance, the Stomiidae (a deep-sea fish family comprising 287 species, of which 41 are present in the 11,638 species dataset), the Salangidae (a family of icefishes with 17 out of the 20 species represented in the 11,638 species dataset), the enigmatic family Regalecidae (with 1 and 2 out of the 3 species represented in our reduced and large datasets, respectively), and the Galaxiidae (even though some benthic species are comprised among the 53 species of this family, of which 27 and 2 are included in the large and the reduced datasets, respectively). The fact that scaleless groups live in open waters supports the hypothesis that scale loss came first, as a likely pre-adaptation to colonize the benthic environment.

While the scaleless phenotype is tightly linked with habitat preference and likely leads to a benthic way of life in ray-finned fishes, the biological meaning of this association is difficult to assert. Some potential explanations could rely in the increased cutaneous respiration in scaleless species (Park and Kom 1999; Park 2002) in a benthic environment characterized by a reduced oxygen content and limited water flow, as compared to open water environments. Another advantage could rely in the increased immune response of scaleless skin (Holm et al. 2017) when confronted to a microbial-rich benthic environment. We could thus argue that fishes having lost their scales are better adapted to the benthic environment, facilitating the colonization of this niche. Because the association between a scaleless phenotype and a benthic way of life has evolved repeatedly and independently many times, these parallel evolutionary trajectories suggest that a scaleless tegument has strong selective value in the benthic environment.

### Regaining scales is unlikely

In our analyses, we inferred very few instances of scale re-acquisition after scale loss events in ray-finned fishes, making the case that scale losses are hardly reversible. The few scale re-acquisition events were inferred either with relatively low probability (e.g. Opisthognathidae family), questioning their validity, or in families with poorly resolved phylogenetic relationships. For instance, in the 304 species phylogeny, we inferred a scale re-acquisition within the Anguilliformes order, more specifically in the Anguillidae family (Fig. S1, Fig. S2 and Fig. S3). However, because the phylogenetic relationships within the Anguilliformes is still debated (e.g. different resolutions of the branching order in the phylogenies of Johnson et al. 2012 and Santini et al. 2013), alternative branching patterns may cancel the inference of a scale re-acquisition. Within Rabosky et al. (2018) phylogeny, scale gains were inferred in relatively enigmatic taxa. The position of such families within this large phylogeny may still lack resolution. Indeed, when looking at lineage-specific phylogenies for the groups in which suspicious scale re-acquisition was inferred, we can observe that the relationships are often different from the ones found in the large phylogeny proposed by Rabosky et al. (2018). As a matter of fact, specific phylogenies of *Clariger and Luciogobius* genera (Yamada et al., 2009), *Parupeneus* (Song et al. 2014), *Lycodes* (Turanov et al. 2017), *Cryptacanthodes* (Radchenko et al. 2011), *Ocosia* (Smith et al. 2018), *Notothenia* (Near et al., 2018), *Lophiocharon* (Arnold and Pietsch, 2012), *Perulibatrachus* (Rice and Bass 2009) and *Stomias* (Kenaley et al. 2014) genera all present differences with the topology of Rabosky et al. (2018) we used in our reconstruction (the problematic subtrees are presented Fig. S11). Consequently, scale re-acquisitions could be artifacts resulting from topological errors.

In one instance (within the Mastacembelidae), a scale gain was found with the Bayestrait method but not with the stochastic mapping, indicating that methodological biases may occur, and that selecting the most appropriate parameter values can be subtle. However, the probability of scale re-acquisition inferred by Bayestrait is not high (<0.75).

In any case, we here show that scale loss events occurred multiple times along the evolutionary history of actinopterygians, while scale re-acquisitions were extremely rare or non-existent. This finding suggests that the gene regulatory network underlying scale formation is difficult to reassemble after it has been dismantled during a scale loss event.

Recent studies showed that the absence of scales in ray-finned fishes may be associated with genetic changes (Liu et al. 2016). For instance, in the secretory calcium-binding phosphoprotein (SCPP) gene family, which is important for scale mineralization in various ray-finned fishes (Liu et al. 2016; Lv et al. 2017), the SCCP1 and SCPP5 genes have been proposed as candidates genes linked to scale presence or absence (Liu et al. 2016). Yet, while these genes are linked to the scaleless phenotypes in some species (e.g. *Ictalurus punctatus, Electrophorus electricus;* Liu et al. 2016), other scale losses could not be linked to these specific genes (e.g. *Sinocyclocheilus anshuiensis;* Lv et al. 2017). As such, different genes and/or set of genes may be underlying the scale presence or absence (Lv et al. 2017) and more research is needed to uncover upstream genetic switches.

### Trunk bony plates emerge on a scaleless tegument

The emergence of TBP occurred in several places of the studied phylogenies (Fig. 1 and Fig. 2 for the 304 and 11,638 species datasets, respectively). The presence of TBP structures is found in different unrelated taxa, and thus bears virtually no phylogenetic signal (Table S4). We here show that there is a common ground needed for the emergence of such plates on the trunk of ray-finned fishes, which is the absence of scales. The mechanistic relations between these two traits remain however uncertain. Interestingly, it also appears that the acquisition of TBP is hardly reversible in the 304 species phylogeny, yet possible but extremely rare in the 11,638 one. This discrepancy can be explained by the fact that the few clades showing TBP losses in the 11,638 species phylogeny are not present in the 304 species dataset (Auchenipteridae, Harpagiferidae, Tetrarogidae), while some clades are present, yet represented with only few species (e.g. Cottidae, Cyclopteridae, Nototheniidae). Interestingly however, most of the TBP losses inferred using the 11,638 species phylogeny occurred in groups which internal phylogeny is not perfectly resolved. The large phylogeny of Rabosky et al. (2018) is indeed different than specialized phylogenies, for instance in the Siluriformes order (Sullivan et al. 2006), the Tetrarogidae family (Smith et al. 2018) and the Nototheniidae and Harpagiferidae families (Near et al. 2018).

The discovery that TBP emerged on taxa with scaleless teguments together with the strong association between scaleless tegument and benthic habitat preference explains the observation that almost all extant fishes displaying TBP have a benthic habitat preference. One main function of scales is the physical protection they provide (Vernerey and Barthelat 2014). It is thus possible that, in the absence of scales, and given the suggested complexity of the genetic control of scale development, simpler alternative developmental pathways can be reached leading to the emergence of a different protective tegument in the form of bony plates on the trunk. The numerous independent acquisitions of a hard armor composed of TBP is indicative of the reduced genetic complexity underlying their emergence given the actinopterygian genetic background. Furthermore, once TBP have been acquired, the low rate of secondary losses indicates that they likely confer some evolutionary advantages, as previously suggested by Vickaryous and Sire 2009.

### Limitations of the study

Our results give new insight into the interconnected evolution among different tegument structures in fish. However, different elements could limit the outcome and interpretation of our study.

First, investigating the evolution of traits that are still debated within the scientific community is a challenging task. Indeed, clear consensus about the distinction between different tegument structures in fish has yet to be reached, and we thus opted for a macro-structural approach differentiating micromeric scale-like elements from macromeric bony plate-like elements. More research is needed to understand the homology among these categories of structures, in particular through paleontological and developmental genetics studies.

Second, we have mentioned some situations that may hamper the complete resolution, and with high confidence, of the ancestral state reconstructions. Errors in the phylogenetic tree and lack of resolution in parts of the tree may mislead the ancestral state reconstructions. Indeed, the robustness of the phylogeny is paramount for proper reconstruction of ancestral states. For instance, the discrepancies between the relationships within the Siluriformes order in the large phylogeny of Rabosky et al. (2018), as compared to the lineage-specific Siluriformes phylogeny of Sullivan et al. 2006 may explain the doubtful loss of TPB we inferred in some Silurifomes taxa. We have also observed that the impressive taxonomic sampling yet coupled with a reduced amount of sequence data characterizing the phylogenetic tree of Rabosky et al. (2018) resulted in some differences when analyzing it with the two ancestral state reconstruction methods (i.e. stochastic mapping vs Bayestrait). To the contrary, the more robust phylogeny of Hughes et al. (2018) but with a reduced taxonomic sampling showed virtually no disparity between the results obtained with the same two methods. In the phylogeny of Rabosky et al. (2018), we pointed controversial phylogenetic relationships within several genera, that is, at a recent phylogenetic scale were high quality sequences of fast evolving markers are required for a fine resolution. For example, some well recognized genera were found to be polyphyletic, a problem that most likely explains the few unexpected recent scale loss events.

Third, the lack of precise morphological knowledge about the traits of interest in some poorly described taxa can lead to erroneous trait attributions, and thus to some artifactual reconstructions (mistaken gains or loss of structures). This situation might be found in the Gobiidae family with the genus *Luciogobius*, in the family Cottidae with the genus *Clinocottus*, or in the family Tetrarogidae with the genus *Ocosia*, among others.

Despite the above-mentioned limitations explaining why our study cannot certify the accuracy of every single reconstructed event along the evolution of the tegument structures in ray-finned fishes, the general patterns we present are robust to changes in analytical methods, dataset size, and phylogenetic uncertainty. Overall, the tested conditions yielded very similar results supporting our conclusions.

### Conclusion

We here demonstrate that scale loss event occurred several times, in an independent manner along the evolution of Actinopterygii, the most species-rich group of vertebrates. We observe that these scale losses are hardly reversible as scale re-acquisition are extremely unlikely. We show that the scaleless phenotype is associated to a benthic habitat preference, and we argue that following a scale loss event, fishes tend to colonize the floors of oceans and water bodies, adopting a benthic lifestyle. The repeated and parallel colonization of the seafloor after a scale loss event indicate that the scaleless phenotype most probably confers a selective advantage in this particular habitat. We also show that the multiple emergences of TBP are phylogenetically independent. We demonstrate that their emergence is dependent over a previous scale loss event. Indeed, these plates are never present in scaled bodies and thus only arise in fish displaying a scaleless tegument, in a “gain after scale loss” evolutionary sequence. The precise mechanisms ruling the interplay between the loss of scales and the emergence of trunk plates remain however to be studied further. Studies focusing on the gene regulatory networks implicated in the transition between tegument structures along evolution could shed light upon the transition between scales and TBP in ray-finned fishes. Finally, our findings support the hypothesis that trunk tegument structures are functional innovations that contributed to the radiation of ray-finned fishes in the aquatic environment.

## Acknowledgments

We thank the University of Geneva BioSC service and Dr. José Manuel Nunes for advices on the statistical tests. We thank Luigi Manuelli and Killian Perrelet for comments on the manuscript and Slim Chraiti for his photographic work. The computations were performed at University of Geneva on the Baobab cluster. This research was supported by the Swiss National Science Foundation (project N° 310030_185327 to JIMB), by the department GENEV, and by the Claraz Donation.

## Author’s contribution

A-L gathered the data and performed the analyses. J.I.M.-B. designed and supervised the study. Both authors wrote and edited the manuscript and approved its submission.

## Data Accessibility

Supplementary material will be available upon acceptance on dryad accession number XXX. They are accessible at the website: https://genev.unige.ch/research/laboratory/Juan-Montoya (under the tab “MORE”).

## Notes

https://genev.unige.ch/research/laboratory/Juan-Montoya

### Competing Interest Statement

The authors have declared no competing interest.

## References

Aberer, A.J., Kobert, K. and Stamatakis, A. 2014. Exabayes: Massively parallel bayesian tree inference for the whole-genome era. Mol. Biol. Evol. 31: 2553–2556.

Alfaro, M.E. 2018. Resolving the ray-finned fish tree of life. Proc. Natl. Acad. Sci. U. S. A. 115: 6107–6109.

Arnold, R.J. and Pietsch, T.W. 2012. Evolutionary history of frogfishes (Teleostei: Lophiiformes: Antennariidae): A molecular approach. Mol. Phylogenet. Evol. 62: 117–129. Elsevier Inc.

Benthon, M.. 2004. Vertebrate Paleontology, 3rd ed. Blackwell Science Ltd.

Covain, R., Fisch-Muller, S., Oliveira, C., Mol, J.H., Montoya-Burgos, J.I. and Dray, S. 2016. Molecular phylogeny of the highly diversified catfish subfamily Loricariinae (Siluriformes, Loricariidae) reveals incongruences with morphological classification. Mol. Phylogenet. Evol. 94: 492–517.

David Johnson, G., Ida, H., Sakaue, J., Sado, T., Asahida, T. and Miya, M. 2012. A “living fossil” eel (Anguilliformes: Protanguillidae, fam. nov.) from an undersea cave in Palau. Proc. R. Soc. B Biol. Sci. 279: 934–943.

Friedman, M. and Brazeau, M.D. 2010. A reappraisal of the origin and basal radiation of the Osteichthyes. J. Vertebr. Paleontol. 30: 36–56.

Fritz, S.A. and Purvis, A. 2010. Selectivity in mammalian extinction risk and threat types: A new measure of phylogenetic signal strength in binary traits. Conserv. Biol. 24: 1042–1051.

Froese, R. and Pauly., D. 2011. Fishbase.

Gemballa, S. and Bartsch, P. 2002. Architecture of the Integument in Lower Teleostomes : Functional Morphology and Evolutionary Implications. 309: 290–309.

Holm, H.J., Skugor, S., Bjelland, A.K., Radunovic, S., Wadsworth, S., Koppang, E.O., et al. 2017. Contrasting expression of immune genes in scaled and scaleless skin of Atlantic salmon infected with young stages of Lepeophtheirus salmonis. Dev. Comp. Immunol. 67: 153–165. Elsevier Ltd.

Huelsenbeck, J.P., Nielsen, R. and Bollback, J.P. 2003. Stochastic mapping of morphological characters. Syst. Biol. 52: 131–158.

Hughes, L.C., Ortí, G., Huang, Y., Sun, Y., Baldwin, C.C., Thompson, A.W., et al. 2018. Comprehensive phylogeny of ray-finned fishes (Actinopterygii) based on transcriptomic and genomic data. Proc. Natl. Acad. Sci. U. S. A. 115: 6249–6254.

Ichiro, S., Mikio, I., Hiroyuki, Y. and Masato, M. 2013. Teeth and ganoid scales in Polypterus and Lepisosteus, the basic actinopterygian fish: An approach to understand the origin of the tooth enamel. J. Oral Biosci. 55: 76–84. Elsevier.

Keating, J.N. and Donoghue, P.C.J. 2016. Histology and affinity of anaspids, and the early evolution of the vertebrate dermal skeleton. Proc. R. Soc. B Biol. Sci. 283.

Kenaley, C.P., Devaney, S.C. and Fjeran, T.T. 2014. The complex evolutionary history of seeing red: Molecular phylogeny and the evolution of an adaptive visual system in deep-sea dragonfishes (stomiiformes: Stomiidae). Evolution (N. Y). 68: 996–1013.

Lees, J., Märss, T., Wilson, M.V.H., Saat, T. and Špilev, H. 2012. The sculpture and morphology of postcranial dermal armor plates and associated bones in gasterosteiforms and syngnathiforms inhabiting Estonian coastal waters. Acta Zool. 93: 422–435.

Liu, Z., Liu, S., Yao, J., Bao, L., Zhang, J., Li, Y., et al. 2016. The channel catfish genome sequence provides insights into the evolution of scale formation in teleosts. Nat. Commun. 7: 11757.

Lv, Y., Kawasaki, K., Li, J., Li, Y., Bian, C., Huang, Y., et al. 2017. A genomic survey of SCPP family genes in fishes provides novel insights into the evolution of fish scales. Int. J. Mol. Sci. 18: 1–11.

Meunier, F.J. and Brito, P.M. 2004. Histology and morphology of the scales in some extinct and extant teleosts. Cybium 28: 225–235.

Mongera, A. and Nüsslein-Volhard, C. 2013. Scales of fish arise from mesoderm. Curr. Biol. 23: R338–R339. Elsevier.

Moyle, P.B. and Cech, J.J. 2004. Fishes, An introduction to Ichthyology, 5th ed. (B. Cummings, ed).

Near, T.J., MacGuigan, D.J., Parker, E., Struthers, C.D., Jones, C.D. and Dornburg, A. 2018. Phylogenetic analysis of Antarctic notothenioids illuminates the utility of RADseq for resolving Cenozoic adaptive radiations. Mol. Phylogenet. Evol. 129: 268–279. Elsevier.

Nelson, J.S., Grande, T.C. and Wilson, M.V.H. 2016. Fishes of the World, 5th ed. Wiley.

Orme, C.D.L., Freckleton, R.P., Thomas, G.H., Petzoldt, T. and Fritz, S.A. 2013. The caper package: comparative analyses of phylogenetics and evolution in R. http://caper.rforge.r-project.org. Google Sch.

Pagel, M., Meade, A. and Barker, D. 2004. Bayesian estimation of ancestral character states on phylogenies. Syst. Biol. 53: 673–684.

Paradis, E., Claude, J. and Strimmer, K. 2004. APE: Analyses of Phylogenetics and Evolution in R language. Bioinformatics 20: 289–290.

Park, J.Y. 2002. Structure of the skin of an air-breathing mudskipper, Periophthalmus magnuspinnatus. J. Fish Biol. 60: 1543–1550.

Park, J.Y. and Kom, I.-S. 1999. Structure and histochemistry of skin of mud loach Misgurnus anguillicaudatus (Pisces, Cobitidae), from Korea. Korean J. ichtyology 11: 109–116.

Porter, M.M., Novitskaya, E., Castro-Ceseña, A.B., Meyers, M.A. and McKittrick, J. 2013. Highly deformable bones: Unusual deformation mechanisms of seahorse armor. Acta Biomater. 9: 6763–6770. Acta Materialia Inc.

Qu, Q., Zhu, M. and Wang, W. 2013. Scales and Dermal Skeletal Histology of an Early Bony Fish Psarolepis romeri and Their Bearing on the Evolution of Rhombic Scales and Hard Tissues. PLoS One 8.

Rabosky, D.L., Chang, J., Title, P.O., Cowman, P.F., Sallan, L., Friedman, M., et al. 2018. An inverse latitudinal gradient in speciation rate for marine fishes. Nature 559: 392–395. Springer US.

Radchenko, O.A., Chereshnev, I.A., Petrovskaya, A. V. and Antonenko, D. V. 2011. Relationships and position of wrymouths of the family cryptacanthodidae in the system of the suborder Zoarcoidei (Pisces, Perciformes). J. Ichthyol. 51: 487–499.

Rambaut, A., Drummond, A.J., Xie, D., Baele, G. and Suchard, M.A. 2018. Posterior summarization in Bayesian phylogenetics using Tracer 1.7. Syst. Biol. 67: 901–904.

Revell, L.J. 2012. phytools: An R package for phylogenetic comparative biology (and other things). Methods Ecol. Evol. 3: 217–223.

Rice, A.N. and Bass, A.H. 2009. Novel vocal repertoire and paired swimbladders of the three-spined toadfish, Batrachomoeus trispinosus: Insights into the diversity of the Batrachoididae. J. Exp. Biol. 212: 1377–1391.

Rivera-Rivera, C.J. and Montoya-Burgos, J.I. 2017. Trunk dental tissue evolved independently from underlying dermal bony plates but is associated with surface bones in living odontode-bearing catfish. Proc. R. Soc. B Biol. Sci. 284.

Santini, F., Kong, X., Sorenson, L., Carnevale, G., Mehta, R.S. and Alfaro, M.E. 2013. A multilocus molecular timescale for the origin and diversification of eels (Order: Anguilliformes). Mol. Phylogenet. Evol. 69: 884–894. Elsevier Inc.

Schultze, H.P. 2018. Hard tissues in fish evolution: History and current issues. Cybium 42: 29–39.

Sire, J.-Y. 1993. Development and fine structures of the bony scutes in corydoras arcuatus (Siluriformes, Callichthyidae). J. Morphol. 215: 225–244.

Sire, J.-Y., Allizard, F., Babiar, O., Bourguignon, J. and Quilhac, A. 1997. Scale development in zebrafish (Danio rerio). J. Anat. 190: 545–561.

Sire, J.Y., Donoghue, P.C.J. and Vickaryous, M.K. 2009. Origin and evolution of the integumentary skeleton in non-tetrapod vertebrates. J. Anat. 214: 409–440.

Sire, J.Y. and Huysseune, A. 2003. Formation of dermal skeletal and dental tissues in fish: A comparative and evolutionary approach. Biol. Rev. Camb. Philos. Soc. 78: 219–249.

Smith, W.L., Everman, E. and Richardson, C. 2018. Phylogeny and Taxonomy of Flatheads, Scorpionfishes, Sea Robins, and Stonefishes (Percomorpha: Scorpaeniformes) and the Evolution of the Lachrymal Saber. Copeia 106: 94–119.

Song, H.Y., Mabuchi, K., Satoh, T.P., Moore, J.A., Yamanoue, Y., Miya, M., et al. 2014. Mitogenomic circumscription of a novel percomorph fish clade mainly comprising “Syngnathoidei” (Teleostei). Gene 542: 146–155. Elsevier B.V.

Stöver, B.C. and Müller, K.F. 2010. TreeGraph 2: Combining and visualizing evidence from different phylogenetic analyses. BMC Bioinformatics 11: 1–9.

Turanov, S. V., Kartavtsev, Y.P., Lee, Y.H. and Jeong, D. 2017. Molecular phylogenetic reconstruction and taxonomic investigation of eelpouts (Cottoidei: Zoarcales) based on Co-1 and Cyt-b mitochondrial genes. Mitochondrial DNA Part A DNA Mapping, Seq. Anal. 28: 547–557.

Vernerey, F.J. and Barthelat, F. 2014. Skin and scales of teleost fish: Simple structure but high performance and multiple functions. J. Mech. Phys. Solids 68: 66–76. Elsevier.

Vickaryous, M.K. and Sire, J.Y. 2009. The integumentary skeleton of tetrapods: Origin, evolution, and development. J. Anat. 214: 441–464.

Wainwright, P.C. and Longo, S.J. 2017. Functional Innovations and the Conquest of the Oceans by Acanthomorph Fishes. Curr. Biol. 27: R550–R557.

Yamada, T., Sugiyama, T., Tamaki, N., Kawakita, A. and Kato, M. 2009. Adaptive radiation of gobies in the interstitial habitats of gravel beaches accompanied by body elongation and excessive vertebral segmentation. BMC Evol. Biol. 9: 1–14.

Zhu, D., Ortega, C.F., Motamedi, R., Szewciw, L., Vernerey, F. and Barthelat, F. 2012. Structure and mechanical performance of a “modern” fish scale. Adv. Eng. Mater. 14: 185–194.

